# A silent killer in the Far North Region of Cameroon: increasing prevalence of hypertension among population living in Kaele

**DOI:** 10.1101/472357

**Authors:** Françoise Raïssa Ntentie, Ousmane Mfopou Mboindi, Gérald Dama, Maxwell Wandji Nguedjo, Boris Ronald Tonou Tchuente, Boris Gabin Kingue Azantsa, Judith Laure Ngondi, Enyong Julius Oben

## Abstract

**Objectives:** Hypertension (HTN) is the major risk factor of cardiovascular diseases. Its prevalence is still in perpetual increase worldwide. The aim of this study was to evaluate the prevalence and risk factors of HTN among Kaele dwellers, in the Far North Region of Cameroon where less attention seems to be paid on awareness and sensitization against overnutrition related diseases.

**Methods:** Two hundred and four participants were recruited during free health campaign on cardiovascular diseases organized from 10-15^th^ February 2017 in kaele. Anthropometric and clinical parameters (weight, height, waist circumference, body mass index, blood pressure and heart rate) were measured. A blood sampling was done for lipid profile analysis. HTN and subtypes were diagnosed according to WHO and IDF definitions respectively meanwhile hypercholesterolemia and hypertriglyceridemia were diagnosed with IDF criteria.

**Results:** The overall prevalence of the HTN was 29.9%. Men were more affected than women (35% vs 22.6%, p<0.05). Participants aged between 50-59 years and > 60 years were more exposed (p<0.05). Forty-one percent (41%) of the hypertensive subjects of the study had systo-diastolic sub-type of HTN meanwhile 31.6% had isolated systolic HTN vs 23% with isolated diastolic HTN. Risk factors associated to HTN were : male gender (OR=2.236; p<0.05); absence of education (OR= 24.296; p<0.05); primary education level (OR=1.933; p<0.05); marital status “married” (OR=3.117; p<0.05), increased age (30-39, 50-59, and > 60 years, respectively with OR=4.113, p<0.05; OR=31.405, p<0.05 and OR=18.694, p<0.05), abdominal obesity (OR= 2.476; p<0.05) and low milky products consumption (OR=2.031, p<0.05).

**Conclusions:** HTN is quite present in Kaele locality and many non-modifiable, modifiable and socioeconomic risk factors significantly contributed to its development.

## Introduction

Non-communicable diseases represent a heavy burden at the global scale and constitute the first cause of death worldwide [1]. According to WHO statistics, those diseases are responsible of 68% of 56 million deaths worldwide. In Africa, there is an important increase of their prevalence and this situation causes, not only a big deficit in the budget of the states or families, but also impairs the health quality of individuals. Amongst these diseases, hypertension (HTN) is the most frequent [2, 3]. Also known as silent killer, HTN causes severe damages on brain, heart, kidney and sometimes premature death when not treated [4]. According to WHO (2011) [5] 30 to 50% of death were attributed to HTN in developing countries. In Cameroon, prevalence of HTN is reported to varied from 31.1% in rural milieu [6], 32.2% in semi-urban [7] to 47.5% in urban milieu [8] with a national average of 31.0% [9]. Despite the efforts being made to reduce the incidence of this silent killer in Cameroon, many research projects initiated on this issue are based mainly in the southern Regions to the detriment of the northern Regions of the Country which seem to face mostly undernutrition related problems such as food insecurity and micronutrient deficiencies [10, 11, 12]. So far, referenced studies carried in the Northern Regions are restricted to headsquares of Region (Ngaoundéré and Maroua) and were done only in the hospital milieu [13, 14, 15, 16]. Thus, there is a need to describe and analyze HTN patterns and its risk factors in order to fill the gap of data and also to contribute in the development of policies for the prevention and management of the pathology and its complications in this part of the country. Therefore, the present study has been initiated and aimed at assessing the prevalence of HTN and identifying the risk factors specific to the population living in Kaele, a locality of the Mayo Kani Division in the Far North Region.

## Material and methods

### Description of study area and population

A cross-sectional and descriptive survey was conducted from 10^th^ to 15^th^ February 2017 in Kaele, head quarter of the Mayo Kani Division in the Far North Region of Cameroon. The sampling was randomized and involved four towns: *Gouzougoui, Kililimbri, Djidoma* and *Lara.* A total of 204 apparently healthy participants aged 18 years and above were recruited during the health campaign organized by the Cameroon Nutritional Science Society on good nutritional practices.

### Ethical considerations

The study protocol was approved by the National Ethics Committee N° 2014/08/488/EC/CNERSH and was conducted in strict compliance with the physical, moral and psychological integrity of all participants; following the principles outlined in the Helsinki Declaration.

### Questionnaire

Data were collected using a questionnaire adapted from WHO STEPwise approach for chronic disease risk factor surveillance-Instrument v2.1. This questionnaire included informations on residence in Kaele (at least one year), smoking, and alcohol. Fruits, vegetables, meat, fish and milk intake were assessed by self-report under the assistance of trained investigators. Alcohol consumption was classified into two categories: abstainers (never consumed) and occasional (drank in the past 12 months) or daily drinkers. Smoking included: manufactured or hand-rolled cigarettes, cigars, smoked, chewed or inhaled products. Participants were classified as abstainers or smokers. Fruits, vegetables, meat, fish, dairy products intake was based on the frequency of intake per week. From 0-1 time per week, intake was classified as low; 2-3 times/week as moderate and 4 to 7 times/week as high. The level of education, marital status, source of income, personal and family history of HTN were also assessed.

### Anthropometric measurements, evaluation of nutritional status and abdominal obesity

Weight, height, waist circumference were measured on participants in light clothing, without shoes and motionless according to standard methods. Body Mass Index (BMI), were computed and categorized according to the World Health Organization (2003) [17] criteria where a BMI <18.5 kg/m^2^ is considered as underweight; BMI range 18,5-25 kg/m^2^ as normal, from 25 to 29 kg/m^2^ individual was considered as overweight, and a BMI ≥ 30 kg/m2 was referred to as obesity.

Abdominal obesity was diagnosed using IDF (2005) [18] criteria (waist circumference ≥80 cm for women and ≥94 cm for men).

### Blood pressure measurement and HTN definition

Blood pressure was measured on seated participant thrice on the right arm at five minutes interval; with uncrossed legs using an Automatic Digital Blood Pressure Arm Monitor SMARTHEART^™^ Mean blood pressure of two closest measures was obtained. Hypertension was diagnosed using two definitions: WHO (1999) [19] (Systolic Blood Pressure (SBP) ≥ 140 mmHg and/or Diastolic Blood Pressure (DBP) ≥ 90 mmHg) and IDF (2005) [18] (SBP ≥ 130 mmHg and/or DBP ≥ 85 mmHg). Hypertension subtypes were referred as per Franklin *et al.* [20] criteria which diagnosis include Isolated Systolic Hypertension (ISH) for a SBP≥130 and DBP<85mmHg; Isolated Diastolic Hypertension (IDH) for a SBP<130 and DBP≥85mmHg and the Systo-Diastolic Hypertension (SDH) with a SBP ≥130 and DBP ≥85mmHg. To classify hypertensive participants according to the grade of illness, the JNC-8 criteria were used as presented in table 1.

**Table 1:** Classification of Blood pressure in Adults (age ≥18 years). (Update JNC-8 Guideline recommandations)

|  | Systolic BP (mmHg) | and/or | Diastolic BP (mmHg) |
| --- | --- | --- | --- |
| **Normal** | $\leq 129$ | | $\leq 84$ |
| **prehypertension** | 130-139 |  | 85-89 |
| **Stage 1 HTN** | 140-159 |  | 90-99 |
| **Stage 2 HTN** | $\geq 160$ | | $\geq 100$ |

### Blood sampling, biochemical analysis and diagnostic of hyperlipidemia

Blood sampling was done in the morning after a 10-hour overnight fast. About 4 ml of venous blood was collected on EDTA tubes by venipuncture in the hand of each participant. The plasma was obtained by centrifugation and aliquots were frozen at −20°C for further biochemical analyses. As biochemical analysis, total cholesterol and triglycerides levels were measured with standard enzymatic spectrophotometric method using ChronoLab Diagnostic Kits in the laboratory. IDF (2006) [21] criteria were used to diagnose hypercholesterolemia (total cholesterol level ≥200mg/dL) and hypertriglyceridemia (triglycerides level ≥150mg/dL).

### Data management and statistical analysis

Data were analyzed using SPSS 16.0 for Windows. Descriptive analysis results were presented as means ± standard deviations for continuous variables and as frequencies for categorial variables. Student t test and chi square test were performed to compare continuous and categorial variables respectively. whereas binary logistic regressions were used to evaluate the relative risk of HTN with statistical significance at p < 0.05.

## Results

### Baseline Characteristics

As shown in Table 2, the study population was constituted of 58.8% male (120) and 41.2 % female (84). However 10.3% percent amongst women were menopausal (n=21). Concerning the marital status, there was 39.7% (n=81) married, 56.4% (115) single and 3.9% (8) widowed or divorced. Participants were mostly from the Moundang ethnic group at 87.7% (179), 3.9% (8) were Toupouri, 3.9% (8) were Guiziga and 4.4% (9) from other minor ethnic groups living in kaele locality. 8.4% (17) were illiterates (no education level), 24.6% (50) had a primary level of education and 67% (137) were up to secondary school. According to nutritional status, 75% (153) participants were normal according to BMI, 14.7% (30) were underweight and only 10.3% (11) were overweight or obese. The observation of food habits and lifestyle revealed a high proportion of alcohol consumption (61.8%; n=126) as well as fruits, vegetables, fish, meat and dairy product consumption.

**Table 2:**
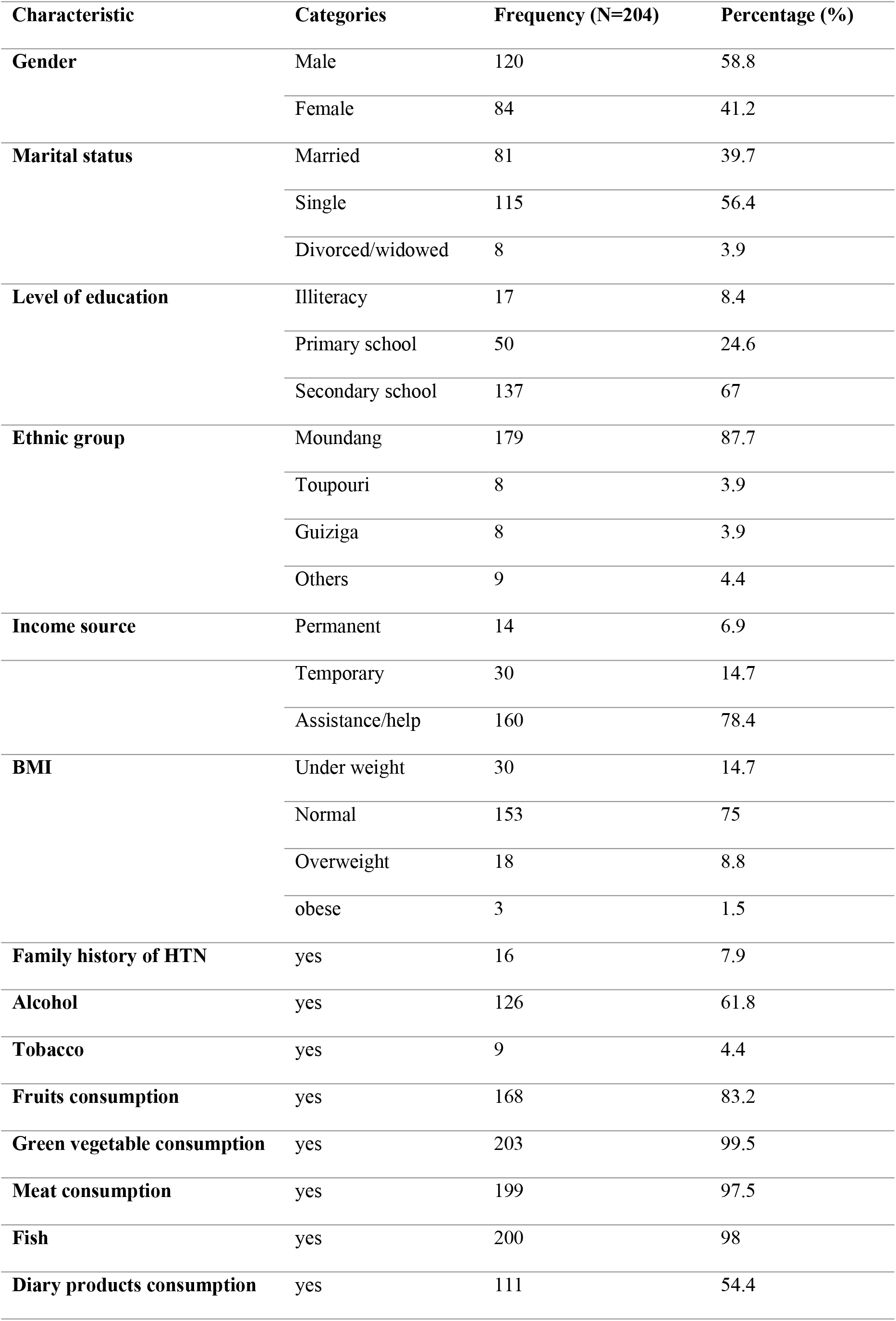
Description of the study population.

| Characteristic | Categories | Frequency (N=204) | Percentage (%) |
| --- | --- | --- | --- |
| **Gender** | Male | 120 | 58.8 |
|  | Female | 84 | 41.2 |
| **Marital status** | Married | 81 | 39.7 |
|  | Single | 115 | 56.4 |
|  | Divorced/widowed | 8 | 3.9 |
| **Level of education** | Illiteracy | 17 | 8.4 |
|  | Primary school | 50 | 24.6 |
|  | Secondary school | 137 | 67 |
| **Ethnic group** | Moundang | 179 | 87.7 |
|  | Toupouri | 8 | 3.9 |
|  | Guiziga | 8 | 3.9 |
|  | Others | 9 | 4.4 |
| **Income source** | Permanent | 14 | 6.9 |
|  | Temporary | 30 | 14.7 |
|  | Assistance/help | 160 | 78.4 |
| **BMI** | Under weight | 30 | 14.7 |
|  | Normal | 153 | 75 |
|  | Overweight | 18 | 8.8 |
|  | obese | 3 | 1.5 |
| **Family history of HTN** | yes | 16 | 7.9 |
| **Alcohol** | yes | 126 | 61.8 |
| **Tobacco** | yes | 9 | 4.4 |
| **Fruits consumption** | yes | 168 | 83.2 |
| **Green vegetable consumption** | yes | 203 | 99.5 |
| **Meat consumption** | yes | 199 | 97.5 |
| **Fish** | yes | 200 | 98 |
| **Diary products consumption** | yes | 111 | 54.4 |

Anthropometric and hemodynamic parameters of the study population (Table 3) revealed that, according to gender, only systolic blood pressure was significantly higher among male (129.78±21.79 mmHg) compared to female (120.64±22.76 mmHg) meanwhile heart rate (p<0.05) was higher among female (83.33±13.44 Pulse/min) than male (75.87±13.14 mmHg).

**Table 3:** Anthropometric and hemodynamic characteristics of the study population.

| Parameters | Population | Female | Male |
| --- | --- | --- | --- |
| Age (years) | $34.16 \pm 17.18$ | $36.02 \pm 17.87$ | $32.85 \pm 16.63$ |
| BMI (kg/m<sup>2</sup>) | $21.40 \pm 3.20$ | $21.79 \pm 3.98$ | $21.12 \pm 2.49$ |
| Waist (cm) | $78.78 \pm 11.25$ | $78.34 \pm 11.62$ | $79.09 \pm 11.02$ |
| SBP (mmHg) | $126.02 \pm 22.60$ | $120.64 \pm 22.76$ | $129.78 \pm 21.79^*$ |
| DBP (mmHg) | $80.82 \pm 12.76$ | $78.80 \pm 11.99$ | $82.22 \pm 13.14$ |
| Heart rate (Pulse/min) | $78.96 \pm 13.74$ | $83.33 \pm 13.44$ | $75.87 \pm 13.14^*$ |

\*significantly different between male and female at $p < 0.05$ with student t test. BMI: body Mass index, *SBP*: systolic blood pressure; *DBP*: diastolic blood pressure.

### Prevalence and characteristic of HTN in Kaele

As presented in table 4, 29.9% (n=61) of participants were hypertensives according to WHO criteria and male were most exposed (35%; n=42) than female (22.6%; n=19) (p<0.05). The results also revealed an important proportion of prehypertensive subjects (18,6%) in the study population. Most hypertensive participants suffering from combine systodiastolic subtype. The overall prevalence also increased with age.

**Table 4:** Prevalence and characteristic of hypertension in the study population.

|  |  | Population % (n) | Female % (n) | Male % (n) |
| --- | --- | --- | --- | --- |
| **Diagnostic criteria** | **WHO** | 29.9 (61) | 22.6 (19)* | 35 (42) |
|  | **IDF** | 48.6 (99) | 36.1 (31)* | 56.7 (68) |
| **HTN stage** | **PreHTN** | 18.6 (38) | 13.4 (12) | 21.7 (26) |
|  | **Stage 1** | 16.2 (33) | 14.3 (12) | 17.5 (21) |
|  | **Stage 2** | 13.8 (28) | 8.4 (7) | 17.5 (21) |
| **Subtypes** | **Isolated diastolic HTN** | 18.2 (18) | 25.8 (8) | 14.7 (10) |
|  | **Isolated systolic HTN** | 34.3 (34) | 32.3 (10) | 35.3 (24) |
|  | **Systo-diastolic HTN** | 47.5 (47) | 41.9 (13) | 50 (34) |
| **Age strata** | **18-29 years** | 30.8 (37/120) | 8.9 (4/45) | 44 (33/75) |
| **% (n/N)** | **30-39 years** | 64.7 (11/17) | 66.7 (4/6) | 63.6 (7/11) |
|  | **40-49 years** | 50 (12/24) | 27.3 (3/11) | 69.2 (9/13) |
|  | **50-59 years** | 93.3 (14 /14) | 90.9 (10/11) | 100 (4/4) |
|  | **≥ 60 years** | 89.3 (25 /28) | 90.9 (10/11) | 88.2 (15/17) |

\*significantly different between male and female at $p < 0.05$ with khi<sup>2</sup> test

### HTN and others cardiometabolic risk factors

The relationship between HTN and others cardiometabolic risk factors such as overweight, abdominal obesity, hypercholesterolemia and hypertriglyceridemia were evaluated (Table 5 and 6). The Pearson correlation between BP and some anthropometric and lipid parameters revealed a positive correlation between BMI and SBP (r=0.139; p=0.048) or DBP (r=0.181; p=0.010); and between waist and SBP (r=0.265; p=0.0001) (Table 5). These observations were also confirmed by the significant higher waist circumference (82.10±10.24 cm; p=0.004) or BMI (21,85±3,66kg/m^2^; p=0.049) among hypertensive group compared to non hypertensive individuals and also by a significantly high prevalence of hypertensive subjects with abdominal fat accumulation (Table 6). For lipid disorders, no significant difference was observed between hypertensive and non hypertensive individuals (Table 6) as well as no relationship between BP level and lipidic parameters (Table 5). But regardless of the prevalence of hyperlipidemia, the results on table 6 shows that the number of subjects with hypercholesterolemia (n=03; 3.3%) and hypertriglyceridemia (n=6; 6.7%) was more higher among hypertensive group compared to non hypertensive individuals.

**Table 5:** Pearson correlation between blood presure level and some anthropometric and lipid parameters.

|  |  | BMI | Waist<br>circumference | Total cholesterol | Triglycerides |
| --- | --- | --- | --- | --- | --- |
| SBP | Correlation coefficient r | **0.139** | **0.265** | 0.137 | 0.054 |
|  | P value | **0.048** | **0.0001** | 0.060 | 0.458 |
| DBP | Correlation coefficient r | 0.131 | **0.181** | 0.066 | 0.043 |
|  | P value | 0.062 | **0.010** | 0.366 | 0.555 |

**Table 6:** Relation between others cardiometabolic risk factors and HTN.

|  | **Population**<br>**(N=190)** | **Hypertensive**<br>**(N=55)** | **Normal**<br>**(N=135)** | **P value** |
| --- | --- | --- | --- | --- |
| Cholesterol (mg/dL) | 116.38±34.46 | 120,62±41,02 | 112,55±26,91 | 0.107 |
| Triglycerides (mg/dL) | 76.42±46.50 | 77,48±45,95 | 75,46±47,20 | 0.765 |
| IMC (kg/m<sup>2</sup>) | 21.40±3.20 | 21,85±3,66 | 20,97±2,63 | **0.049** |
| Waist circumference (cm) | **78.78±11.25** | 81,33±10,04 | 76,43±11,84 | **0.004** |
| Hypercholesterolemia<br>(Chol≥200mg/dL) (% (n)) | 1.6 (3) | 3.3 (3) | 0 (0) | 0.066 |
| Hypertriglyceridemia<br>(TAG≥150mg/dL) (% (n)) | 4.2 (8) | 6.7 (6) | 2.0 (2) | 0.110 |
| Overweight<br>(BMI≥25kg/m<sup>2</sup>) (% (n)) | 10,3 (21) | 13,1 (13) | 7,6 (8) | 0,195 |
| Abdominal fat distribution<br>(% (n)) | 23.1 (40) | 31.3 (26) | 15.6 (14) | 0,014 |

Chol : Cholesterol ; TAG : Triglycerides. Results are expressed as means±Standard deviation for lipid blood level, IMC and Waist and as % (n) for lipidemia.

### Influence of some risk factor on the incidence of high blood pressure in the study population

Table 7 shows gender influence on occurrence of elevated blood pressure. It appears that men were 2.2 times more exposed to elevated blood pressure than women. Regarding the level of education, those with no education were 24.3 times more likely to be at risk of high blood pressure (HBP), and those with primary education level were 1.9 times more likely to suffer hypertension than those with a secondary education level. For marital status, “married” individuals increased by 3.1 times the risk of HTA compared to “single” individuals. With respect to age, individuals aged 30-39, 50-59, 60 and older were respectively 4.1 times; 31.4 times and 18.7 times more exposed to high blood pressure than the younger ones. Regarding the consumption of dairy products, those who did not consume them were 2 times more exposed to hypertension. Another important fact was the increased risk of HTN (2.4 times) with the presence of abdominal fat distribution (p<0.05).

**Table 7:** Odd ratio of elevated blood pressure according to some modifiable and non-modifiable risk factors.

|  |  | **Odd ratio (CI 95%)** | **p value** |
| --- | --- | --- | --- |
| **Gender** | Female | 1 |  |
|  | **Male** | **2.236 (1.262-3.960)** | **0.006** |
| **Education level** | Secondary | 1 |  |
|  | **illiteracy** | **24.296 (3.130-188.595)** | **0.002** |
|  | **Primary** | **1.933 (1.003-3.723)** | **0.049** |
| **Marital status** | Single | 1 |  |
|  | **Married** | **3.117 (1.724-5.633)** | **0.0001** |
|  | Widowed/divorced | 2.897 (0.659-12.736) | 0.159 |
| **Age (years)** | 18-29 | 1 |  |
|  | **30-39** | **4.113 (1.414-11.960)** | **0.009** |
|  | 40-49 | 2.243 (0.922-5.457) | 0.075 |
|  | **50-59** | **31.405 (3.981-247.745)** | **0.001** |
|  | **60 and above** | **18.694 (5.310-65.816)** | **0.0001** |
| **Income source** | Permanent | 1 |  |
|  | Temporary | 1.006 (0.324-3.124) | 0.992 |
|  | Assistance/help | 0.466 (0.149-1.452) | 0.188 |
| **overweight** | no | 1 |  |
|  | yes | 1.833 (0.725-4.633) | 0.200 |
| **Abdominal fat** | no | 1 |  |
| **accumulation** | yes | **2.476 (1.187-5.164)** | **0.016** |
| **Alcohol** | no | 1 |  |
| **consumption** | yes | 1.632 (0.922-2.889) | 0.093 |
| **Tobacco use** | no | 1 |  |
|  | yes | 3.918 (0.794-19.339) | 0.094 |
| **Fruits consumption** | yes | 1 |  |
|  | no | 0.717 (0.340-1.513) | 0.382 |
| **Vegetables** | yes | 1 |  |
| **consumption** | no | / | / |
| **Meat consumption** | no | 1 |  |
|  | yes | 0.701 (0.115-4.286) | 0.701 |
| **Fish consumption** | yes | 1 |  |
|  | no | 0.347 (0.035-3.392) | 0.363 |
| **Dairy products** | yes | 1 |  |
| **consumption** | no | **2.031 (1.160-3.554)** | **0.013** |

## Discussion

The prevalence of hypertension observed in Kaele locality (29.9%) was lower than that 31.0% noted by Kingue *et al.* [9] in Cameroon. This difference could be explained by the fact that Kaele is a less urbanized area and studies shown that urbanization increases the risk of chronic diseases [22]. In the general population, the systo-diastolic subtype was 41% followed by isolated systolic hypertension subtype (31.6%). In women, isolated systolic hypertension was the dominant subtype (47.4%), while among men, systo-diastolic hypertension was the most prevalent (42.9%). The results obtained was different to those of Azantsa *et al.* [23, 24] who noted that the isolated diastolic subtype was more common in Cameroonian obese and metropolis dwellers, respectively. The lower percentage of isolated diastolic HTA obtained in our study (23%) compared to that obtained by Azantsa *et al.* [23] (32.8%), would be attributed to the lower degree of urbanization of the city of Kaele compared to the metropolis Yaounde. In fact, the lifestyle of urban population are risk factors for high blood pressure (sedentary lifestyle, high consumption of alcohol …) [25]. In addition, this isolated diastolic subtype is strongly correlated with obesity; which was not the case among Kaele dwellers with only 10.3% overweight (Table 6). The significant difference between male and female HTA found in this study (Table 4) was consistent with most published data in Africa and elsewhere in the world [26, 27, 28,]. However, it should be noted that in some countries, the prevalence has been higher among women [29]. Indeed, hypertension is more common in men before age 50, thereafter, the trend is reversed [30]. In fact, ovarian hormones, especially estrogens [31], appear to play an important protective role through the modulation of the renin-angiotensin-aldosterone system [32], an effect on cardiovascular function via their kidneys, heart, vascular system and even the central nervous system [33]. Women would therefore be much more protected against hypertension before menopause [34]. Only 10.2% of the women in this study were menopausal, hence the higher prevalence among men.

Exploration of relationship between high blood pressure and some biochemical parameters (table 5–6) revealed no significant difference of lipid profile between normal participants and those with elevated blood pressure meaning that HTN is not a matter of lipid abnormalies in this population, but the presence of such situation can contribute to increase the incidence of HTN among Kaele inhabitants according to mechanisms described by many studies [35, 36].

In fact, atherogenic lipid abnormalities clearly cause endothelial dysfunction possibly through impaired nitric oxide production and activity, as well as alterations in endothelin-1, endothelin A and B receptor expression the endothelial dysfunction could lead to an inability or difficulty in vasodilatation to appropriate stimuli and eventually to increased resting BP [37].

An important finding of the present study was the high risk of elevated blood pressure among subjects with abdominal fat accumulation (OR= 2.031; IC : 1.160-3.554) (table 7). Knowing that the general prevalence of overweight was low (10.3%) (Table 5), and that kaele city is a less urbanized area calling to become more urbanized, one can be afraid by the increased prevalence of weight gain and later hypertension in this population in the upcoming years if nothing is done. Moreover, the results also revealed an important number of prehypertensive participants (Table 4), although it is not yet the pathologic stage of the disease, it permits to identify those who are likely to progress to stage 1, and needs preventive measurements to reduce the expansion of the disease [38]. As such, the identification of specific risk factors for a population become important and the analysis of data (Table 7) shows some specificities for kaele dwellers. Regarding the level of education, those with no education were 24.3 times higher than those with a primary level 1.9 times more at risk of hypertension than those with a secondary level. These results corroborated those of Bovet *et al.* [39] who also found higher rates of hypertensive subjects with low levels of education. Indeed, schooling would promote a better knowledge of the disease and the means to avoid it. Thus, the establishment of education and awareness programs could be a good mean of prevention. Marital status “married” increased the risk of hypertension by 3.1 times compared to “single” status. This result was contrary to that of Hawkley *et al* [40] who showed a direct link between loneliness and systolic blood pressure. In addition, some studies [41, 42] have shown that married people are less likely to be hypertensive than single people. However, a study conducted in South Africa showed that living in a marriage was negatively associated with hypertension. It appears that among unmarried persons, a reduction in the rating of hypertension (OR = 0.30, p <0.05) was noted compared to the situation of married couples; while widows were associated with an increase in HTA odds (OR = 2.43, p <0.05). This could be due to the stress experienced by some married people since the association between stress and hypertension has been proven. In fact, stess leads to a chronic increase in the secretion of catecholamine and cortisol resulting in a state of insulin resistance, visceral obesity, high levels of triglycerides and low levels of HDL-cholesterol associated with hypertension [43]. In terms of age, individuals aged 30-39, 50-59 and 60 and older were respectively 4.1 times; 31.4 times and 18.7 times more exposed to high blood pressure than the younger ones (Table 7). These results were consistent with several prospective and observational surveys, which found a positive relationship between age and high blood pressure in a number of populations, regardless geographic, cultural and socioeconomic characteristics as reported by Guimaraes [44], who noted that among 25 to 34 years old, the rate of hypertension ranged from 5.5 to 17.8%, while among those aged 55 to 59 this rate rose to 41%. The reduction of elasticity (increased rigidity) of large arteries is pointed out [45] leading to endothelial dysfunction that develops over time and hence contributes to increased arterial stiffness in the elderly with isolated systolic hypertension [37].

No parameter of the food behavior investigated in this study significantly explained the occurrence of hypertension in this population except the consumption of dairy products; those who did not eat were twice as likely to be exposed (Table 7). Indeed, calcium is one of the numerous nutrients responsible for the beneficial effect of dairy products on the control of blood pressure [46]. Other minerals such as magnesium and potassium may also help regulate blood pressure, but their individual contributions are hard to detect when present in calcium-rich foods [47]. The most important factor could be attributed to bioactive peptide like casein, whose inhibitory effect on the angiotensin-1 converting enzyme has already been demonstrated in the process of controlling blood pressure. Other studies have revealed that certain milk peptide derivatives also have hypotensive effects via endothelin 1 modulation, performed by the endothelial cell [48].

## Conclusion

The present study suggests that Kaele populations were prone to hypertension with a predominance of the systodiastolic subtype. In addition, older age, male gender, abdominal fat accumulation, “married” individuals, low educational attainment, and low consumption of dairy products were the main risk factors for hypertension amongst population of this locality. As with undernutrition or nutrient deficiency problem, the implementation of programs against overnutrition related diseases becomes a need in the northern part of country. This will help to prevent the expansion of cardiovascular risk factors like hypertension the silent killer and then to solve the problem of double burden of malnutrition.

## Study limitations

A few number of participants and the absence of data related to salt consumption of these Kaele dwellers constituted the limitations of our study.

## Author Contributions

Study conception and design: NFR. Data collection and entering: FRN, MWN, BRTT, GD, OMM. Biochemical analysis : MWN, BRTT, OMM. Statistical analysis and interpretation: FRN, BGKA. Drafting FRN. Manuscript revision: JLN, EJO. All the authors approved the final version of the manuscript.

